# Automated lesion segmentation with BIANCA: impact of population-level features, classification algorithm and locally adaptive thresholding

**DOI:** 10.1101/437608

**Authors:** Vaanathi Sundaresan, Giovanna Zamboni, Campbell Le Heron, Peter M. Rothwell, Masud Husain, Marco Battaglini, Nicola De Stefano, Mark Jenkinson, Ludovica Griffanti

## Abstract

White matter hyperintensities (WMH) or white matter lesions exhibit high variability in their characteristics both at population- and subject-level, making their detection a challenging task. Population-level factors such as age, vascular risk factors and neurodegenerative diseases affect lesion load and spatial distribution. At the individual level, WMH vary in contrast, amount and distribution in different white matter regions.

In this work, we aimed to improve BIANCA, the FSL tool for WMH segmentation, in order to better deal with these sources of variability. We worked on two stages of BIANCA by improving the lesion probability map estimation (classification stage) and making the lesion probability map thresholding stage automated and adaptive to local lesion probabilities. Firstly, in order to take into account the effect of population-level factors, we included population-level lesion probabilities, modelled with respect to a parametric factor (e.g. age), in the classification stage. Secondly, we tested BIANCA performance when using four alternative classifiers commonly used in the literature, with respect to K-nearest neighbour algorithm currently used for lesion probability map estimation in BIANCA. Finally, we propose LOCally Adaptive Threshold Estimation (LOCATE), a supervised method for determining optimal local thresholds to apply to the estimated lesion probability map, as an alternative option to global thresholding (i.e. applying the same threshold to the entire lesion probability map). For these experiments we used data from a neurodegenerative cohort and a vascular cohort.

We observed that including population-level parametric lesion probabilities with respect to age and using alternative machine learning techniques provided negligible improvement. However, LOCATE provided a substantial improvement in the lesion segmentation performance when compared to the global thresholding currently used in BIANCA. We further validated LOCATE on a cohort of CADASIL (Cerebral autosomal dominant arteriopathy with subcortical infarcts and leukoencephalopathy) patients, a genetic form of cerebral small vessel disease characterised by extensive WMH burden, and healthy controls showing that LOCATE adapts well to wide variations in lesion load and spatial distribution.

## 1. Introduction

White matter hyperintensities (WMH), or white matter lesions, are common radiological abnormalities often associated with cognitive impairment, and one of the main signs of cerebral small vessel disease (SVD) (Pantoni, 2010). However, despite their clinical importance, accurate automated detection of white matter hyperintensities of presumed vascular origin (WMH, also known as white matter lesions, Wardlaw et al., 2013) is very challenging due to the high variability of WMH characteristics both between-and within-subjects. For example, at the population-level, the amount and distribution of white matter lesions have been associated with various factors such as cognition, vascular risk factors and neurodegenerative diseases (Li et al., 2013; Debette and Markus, 2010). WMH also occur more commonly at older age (Simoni et al., 2012; Vannorsdall et al., 2009). At the subject-level, generally WMH exhibit spatial heterogeneity and do not appear with the same contrast in all regions of the brain on structural MRI images (Hernández et al., 2017). For example, typically, periventricular WMH are brighter on T2-weighted, FLAIR or proton density images and bigger compared to deep ones (Kim et al., 2008), and hence can be detected more easily.

The variability in lesion characteristics can affect the performance of supervised lesion segmentation methods at different stages. The estimation of the probability for each voxel to be a lesion (lesion probability map) is affected by population-level factors (such as age, cognition, vascular risk factors and pathological conditions) (Rostrup et al., 2012; Zamboni et al., 2017) and lesion load (Dyrby et al., 2008; Anbeek et al., 2004). Once a lesion probability map is generated, the determination of the threshold used to obtain binary maps is affected by lesion load and the variation in local intensities. In fact, heterogeneity in signal intensity affects lesion probability values determined by segmentation algorithms (De Boer et al., 2009; Anbeek et al., 2004; Dyrby et al., 2008), since brighter lesions (e.g. periventricular WMH) are typically assigned higher probabilities than those with lower contrast (e.g. deep WMH). This makes it difficult to determine the optimal threshold for obtaining the final binary lesion maps. In fact, applying a global threshold (i.e. the same threshold to all the voxels of the lesion probability map) typically results either in the exclusion of deep lesions or overestimation of periventricular lesions. This often results in trading off either sensitivity or specificity of lesion detection (Anbeek et al., 2004).

In this work, our objective was to improve our recently developed WMH segmentation method Brain Intensity AbNormality Classification Algorithm (BIANCA, Griffanti et al., 2016) to overcome population- and subject-level variability in the lesion characteristics. We aim to achieve this by systematically exploring ways of improving various stages of the lesion probability map estimation, and also by developing a method for thresholding that allows for spatial variability of lesion probability map.

Regarding the estimation of lesion probability map, we tested the inclusion of the population-level lesion probabilities with respect to a parametric factor (in our case age) and explored the performance of alternative machine learning methods.

One way to include population-level probabilities is to obtain the distribution of lesions within a population with respect to a parametric factor and use it in BIANCA as either spatial prior or additional feature in the WMH segmentation. In this work, we modelled the spatial distribution of lesions within a population with respect to age to obtain a parametric lesion probability map as described in our previous study (Sundaresan et al., 2018, *Preprint, under review*).

Regarding the alternative machine learning techniques, we explored four other supervised classifiers used in the existing literature and compared their lesion segmentation results with those obtained from the existing K-nearest neighbour (KNN) classification algorithm in BIANCA. Neural networks (Dyrby et al., 2008), support vector machines and adaboost classifier (Wen and Sachdev, 2004) have been already used for WMH segmentation, while random forests have been used for the segmentation of various structures and pathological signs that have similar characteristics as WMH on structural MR images (Geremia et al., 2011; Mitra et al., 2014; Yamamoto et al., 2010).

Aiming to improve the thresholding stage, we propose LOCATE (LOCally Adaptive Thresholds Estimation), a supervised method to determine optimal local thresholds for binarising the subject-level lesion probability map to take into account the spatial heterogeneity in lesion probabilities. LOCATE is guided by local probability values and exploits different lesion characteristics to determine local thresholds. In this work we present LOCATE with specific applicability to BIANCA, however, in principle LOCATE can be applied to the lesion probability map obtained by any method, provided the availability of a training data with manual lesion masks. We evaluated LOCATE on two clinical datasets: a neurodegenerative cohort and a vascular cohort, and later validated it on two additional datasets with very high and negligible lesion loads to further test its robustness.

## 2. Materials and methods

### 2.1 Datasets

In this work we used four datasets to study the effect of variations in lesion characteristics on the WMH segmentation. The datasets are diverse in terms of population, and therefore WMH load and characteristics. They were acquired with a similar acquisition protocol on two different scanners (details are provided below).

#### 2.1.1 Dataset 1: Neurodegenerative cohort – Oxford Project to Investigate Memory and Ageing (OPTIMA)

This is a subset of the same dataset we used in our previous work (Griffanti et al., 2016) to optimise BIANCA (21 subjects with manual lesion segmentation available – lesion load range: 1878 – 89259 mm^3^). Briefly, the dataset includes MRI data from 9 subjects with probable Alzheimer’s Disease, 5 with amnestic mild cognitive impairment and 7 cognitively healthy control subjects recruited from the OPTIMA study and from the Memory Assessment Clinic at the John Radcliffe Hospital in Oxford (Zamboni et al., 2013) (age range 63 – 86 years, mean age 77.1 ± 5.8 years, F:M = 10:11). MRI images were acquired at the University of Oxford OCMR centre on a 3 T Siemens Trio scanner using a T2-weighted, fluid-attenuated inversion recovery (FLAIR) research sequence (TR/TE = 9000/89 ms, flip angle 150°, FOV 220 mm, voxel size 1.1 × 0.9 × 3 mm) and a high-resolution T1-weighted images (3D MP-RAGE) were also acquired (TR/TE = 2040/4.7 ms, flip angle 8°, FOV 192 mm, voxel size 1 mm isotropic) (see Griffanti et al., 2016 for more details).

#### 2.1.2 Dataset 2: Vascular cohort – Oxford Vascular Study (OXVASC)

The dataset consists of 18 consecutive eligible participants in the OXVASC study (Rothwell et al., 2004), who had recently experienced a minor non-disabling stroke or transient ischemic attack (age range 50 – 91 years, mean age 73.27 ± 12.32 years, F:M = 7:11). Scanning was performed at the Oxford Acute Vascular Imaging Centre (AVIC) on a 3 T Siemens Verio scanner using a T2-weighted, FLAIR sequence (TR/TE = 9000/88 ms, flip angle 150°, FOV 192 mm, matrix size 174 × 52 × 192, voxel size 1 × 3 × 1 mm) and a high resolution T1-weighted sequence (3D MP-RAGE sequence, TR/TE = 2000/1.94 ms, flip angle 8°, FOV 256 mm, matrix size 208 × 256 × 256, voxel size 1mm isotropic), and diffusion-weighted imaging (TR/TE= 8000/86 ms, GRAPPA factor 2, flip angle 16°, FOV 192 mm, voxel-size 2×2×2 mm, 32 directions, b-value 1500 s/mm^2^). Manual lesion segmentation was available for all 18 images (lesion load range: 3530 – 83391 mm^3^).

#### 2.1.3 Dataset 3: Cerebral autosomal dominant arteriopathy with subcortical infarcts and leukoencephalopathy (CADASIL) cohort

The dataset consists of 15 patients with CADASIL (age range 33 – 70 years, mean age 53.73 ± 11.31 years, with female to male ratio, F:M = 11:4) (Le Heron et al., 2018, in press). CADASIL is a genetic form of small vessel disease caused by mutations within the NOTCH-3 gene. It is the most common heritable cause of stroke and vascular dementia in adults and it is characterised by extensive damage to white matter brain regions (Chabriat et al., 2009). These patients are younger than those with sporadic SVD, without confounding factors of co-existent neurodegenerative pathology (Attems and Jellinger, 2014). They therefore provide a model of pure SVD in which to investigate WMH (Charlton et al., 2006). Scanning was performed using the same scanner and acquisition parameters as the OXVASC dataset. Manual segmentation was not available for this dataset.

#### 2.1.4 Dataset 4: Healthy controls (HC)

This dataset consists of 19 healthy controls, age-matched with the CADASIL subjects (described in Dataset 3) with age range 29 – 70 years, mean age 54.58 ± 11.25 years, F:M = 6:13. Scanning was performed using the same scanner and acquisition parameters as datasets 2 and 3. Manual segmentation was not available for this dataset.

### 2.2 Use of population-level parametric lesion probability map (PPLPM)

Population-level parametric lesion probability maps (PPLPMs) describe the pattern of lesion distribution with respect to a specific parametric factor. Therefore, they could provide useful additional information to improve lesion segmentation. For our experiments in this work, we considered age as our parametric factor of interest, since relationship has been established between age and WMH distribution (Simoni et al., 2012; Vannorsdall et al., 2009).

We modelled the PPLPM with respect to age within a population consisting of 474 subjects, as described in our previous work (Sundaresan et al., 2018, *Preprint, under review*). Briefly, our Bayesian spline model takes binary lesion maps of individual subjects of the population as inputs and generates a 4D parametric lesion probability map, with age (grouped at intervals of 3 years) along the 4^*th*^ dimension. The resulting parametric lesion probability map indicates the probability of lesion occurrence at a specific age group at each voxel. Figure 1 shows the PPLPM at two example age groups, corresponding to 29-31 years and 59-61 years.

**Figure 1:**
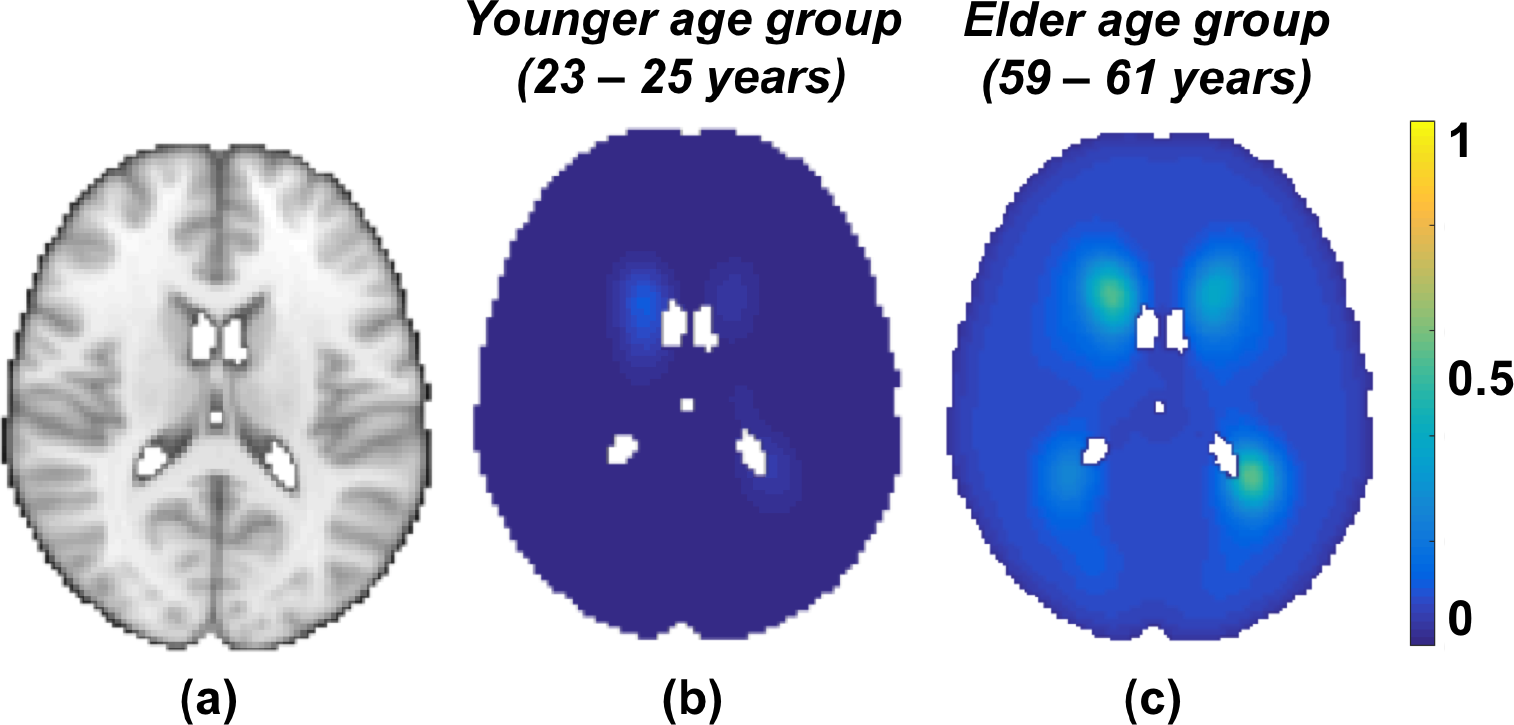
Parametric lesion probability maps at two example points in the parametric dimension, corresponding to two age groups (all the images are shown at z = 45 in MNI space, (a); in the younger age group (29 – 31 years, b) the lesion probability is very low throughout the brain, while in the older age group (59 – 61 years, c) the lesion probability is higher, especially in the periventricular regions.

We used the PPLPM in BIANCA in two ways: either as an additional feature to the KNN classifier, or post-multiplying the PPLPM with the subject’s lesion probability map that is obtained using BIANCA with existing features.

For the first experiment, we implemented the PPLPM in BIANCA as follows: (1) we included the PPLPM (4D volume in the MNI space with 4^*th*^ dimension representing different age groups) as an additional input, (2) we included age as extra input in BIANCA and used it to select the appropriate 3D map from PPLPM corresponding to the specific age group and (3) we transformed the 3D map from the MNI space to the subject’s native space before extracting its probability values as features to be used by the KNN classifier. For the second experiment, we multiplied the age-specific 3D map from PPLPM, transformed to the subject’s native space, with lesion probability map obtained from BIANCA (with the existing options and no additional features). We also tested the effect of using an average 3D map across all age groups instead of the age-specific one.

We performed both experiments on the OPTIMA dataset (keeping all the other options constant, as optimised in (Griffanti et al., 2016)), and evaluated the performance with respect to the manual masks as described in section 2.5.

### 2.3 Comparison of alternative classifiers within BIANCA

Currently, BIANCA is based on the KNN algorithm. In this work we assessed the performance of four other classifiers: random forest (RF), neural networks (NN), support vector machine (SVM) and adaboost (AB). First we optimised the parameters for each classifier based on their area under the curve values from ROC curves, and then compared their results with KNN for the existing features. Table – 1 provides the list of available parameters (from the python *scikit-learn* package) and the parameters that were considered for tuning in the optimisation step.

For initial parameter tuning, we selected four subjects from the OPTIMA dataset with four different lesion loads: low, medium, high and very high (ranging from 5409 – 89259 mm^3^), see figure 2). After optimisation, we applied the four classifiers on the remaining 17 subjects from the OPTIMA dataset (keeping all the other options constant, as optimised in Griffanti et al., 2016) and evaluated the performance with respect to the manual masks as specified in section 2.5.

**Figure 2:**
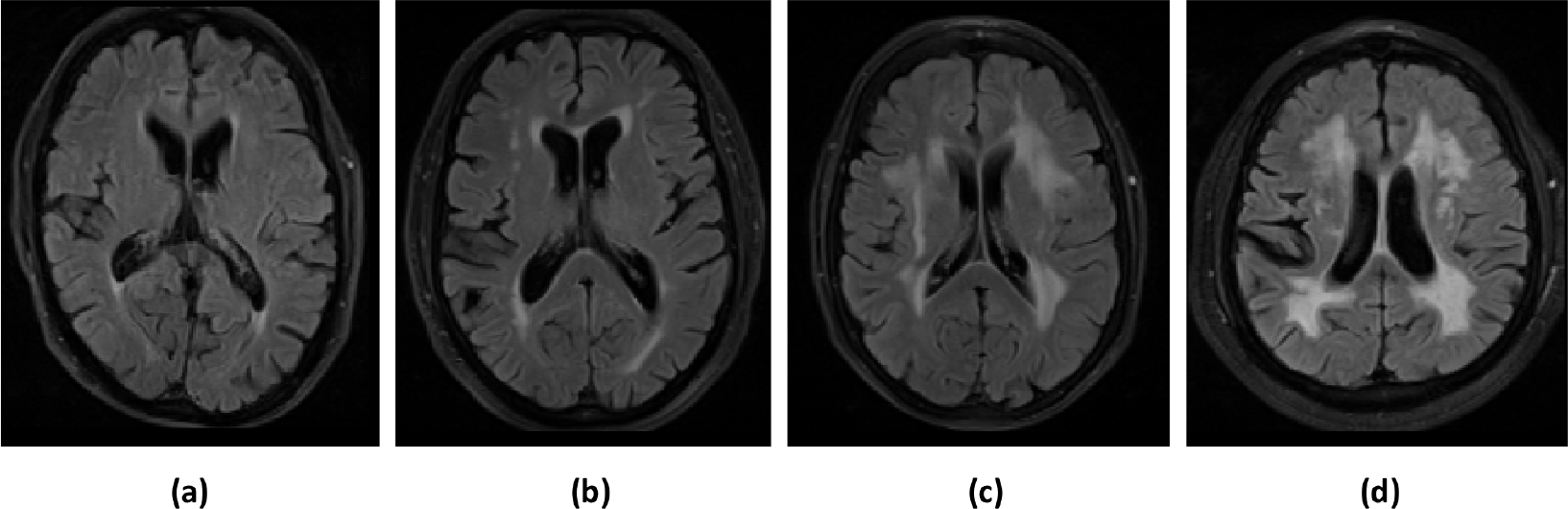
Examples of subjects with different WMH lesion loads used for parameter tuning of the classifiers: (a) low, (b) medium, (c) high and (d) very high lesion load. The manually segmented lesion volumes for low, medium, high and very high lesion loads are 5409 mm^3^, 21005 mm^3^, 50585 mm^3^ and 89259 mm^3^ respectively.

### 2.4 LOCally Adaptive Threshold Estimation (LOCATE)

In order to overcome the impact of spatial heterogeneity of lesion probabilities due to changes in lesion contrast, load and distribution on the final thresholded WMH map, we propose a method that determines spatially adaptive thresholds at different regions of the lesion probability map. LOCATE (LOCally Adaptive Threshold Estimation) takes as input the subject-level lesion probability map obtained from a lesion segmentation algorithm (in our case, BIANCA) and estimates local thresholds in two steps by dividing the lesion probability map into sub-regions (using Voronoi tessellation), extracting local characteristics (features) within those sub-regions and estimating the optimal local threshold values based on the extracted features using a supervised learning method.

#### 2.4.1 Voronoi tessellation and feature extraction

Firstly, we detected local maxima points *M*_*i*_, where *i* = 1…*N* on the lesion probability map to identify the plausible lesion locations (indicated by red dots in figure 3b). In order to avoid spurious local maxima points due to isolated voxels, we spatially smoothed the lesion probability map with a Gaussian kernel of standard deviation, *σ* = 0.5, prior to local maxima detection.

**Figure 3:**
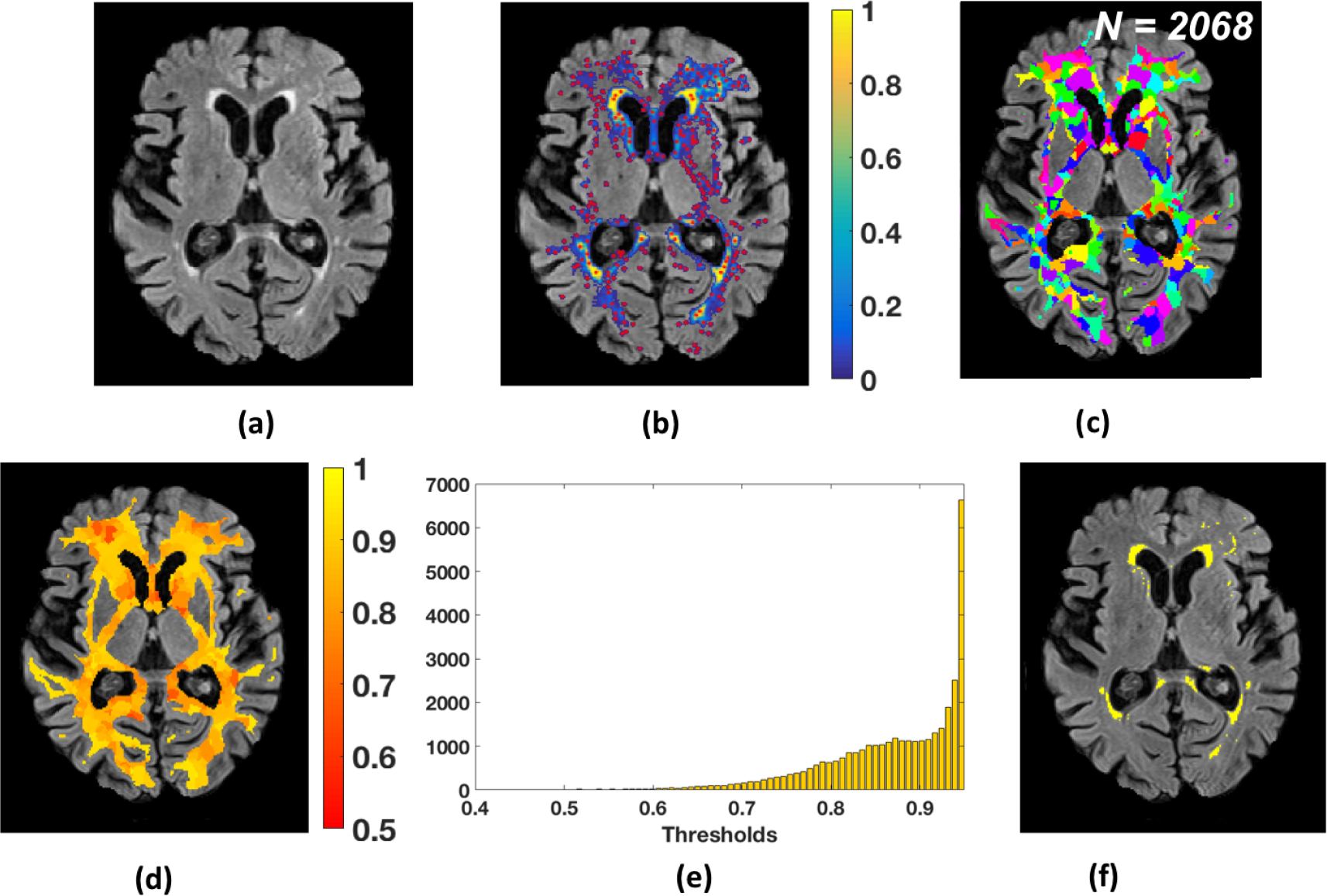
LOCally Adaptive Threshold Estimation (LOCATE). (a) FLAIR image; (b) detection of local maxima (*M*_*i*_, red dots) on the smoothed lesion probability map (blue-yellow); (c) Voronoi spaces *V*_*i*_ around local maxima points (indicated by random colours); (d) threshold map showing the local threshold values obtained from the regression model; (e) histogram of thresholds *Th opt*; (f) final binary lesion map obtained by applying the thresholds (d) on the lesion probability map (b).

We then tessellated the lesion probability map based on local maxima *M*_*i*_ into *N* Voronoi polygons *V*_*i*_, (figure 3b) around the maxima *M*_*i*_. In order to ensure that our Voronoi polygons are within the region of interest (in our case, the brain white matter), we constrain the Voronoi polygons by a white matter mask obtained from FSL FAST tissue segmentation as described in Griffanti et al., 2016.

Within each Voronoi polygon *V*_*i*_, we applied different levels of thresholds (*Th*) from 0 to 0.9 with incremental steps of 0.05, and extracted the following features at each *Th*:

1. Mean greyscale intensity of the image used to identify the lesions (in our case, FLAIR) within the thresholded region
2. Distance between the ventricles and the center of gravity of the thresholded region. Lateral ventricles were were segmented from T1 images as described in Griffanti et al., 2017 and the distance from the ventricle mask was calculated using the FSL command distancemap).
3. Volume of the thresholded region

Due to lack of a simple relationship between the features and the thresholds (figure 4), we determined the optimal local threshold for each *V*_*i*_ using a random forest (Breiman, 2001) regression model with 1000 trees and min leaf size of 5.

**Figure 4:**
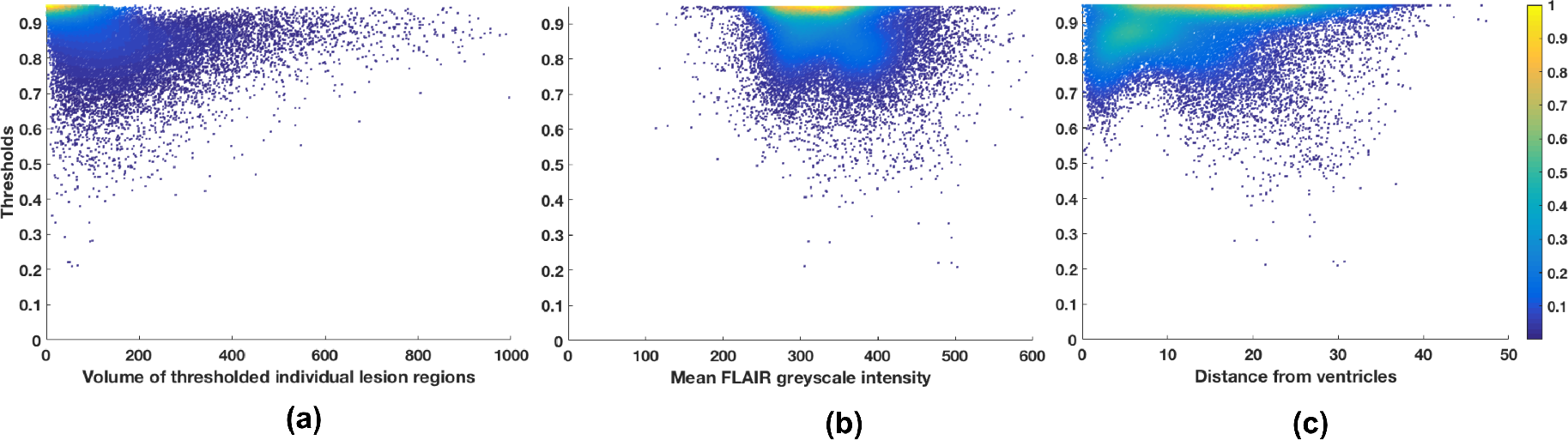
Features extracted from each Voronoi polygon plotted against the threshold values for 21 subjects from the OPTIMA dataset: (a) volume of individual lesion region, (b) mean greyscale intensity and (c) distance of the centre of gravity from the ventricles mask. A simple straightforward relationship was not found between the features and the thresholds, indicating the need for a machine learning based method.

*Training phase*: For each *V*_*i*_, among the set of thresholds *Th* we determined the highest value *Th*_*max*_ at which we obtained the best similarity index with respect to the manually segmented binary lesion mask. We then trained the random forest regression model with the above features against *Th*_*max*_.

*Testing phase*: We applied the trained regression model to get the optimal threshold *Th_opt* for each *V*_*i*_ in the test images. Figure 3d shows an instance of *Th_opt* map. Note that the periventricular region shows higher threshold values compared to the deep region, indicating the variation in lesion probabilities. Moreover, the histogram of thresholds *Th_opt* (figure 3e) obtained for an individual image shows a wider range of local threshold values depending on the local lesion characteristics and spatial distribution. As a final step, we thresholded the lesion probability map within each *V*_*i*_ using the corresponding *Th_opt* to get the final binary lesion map (figure 3f).

#### 2.4.2 LOCATE evaluation and validation

We initially evaluated LOCATE on OPTIMA and OXVASC datasets. For the OPTIMA dataset, the lesion probability maps used as input were obtained using BIANCA with the parameters specified in Griffanti et al., 2016. For the OXVASC dataset, the lesion probability maps were obtained using BIANCA with intensity features from FLAIR, T1 and mean diffusivity images (additional options used: local average intensity within a kernel of size = 3 voxels; MNI coordinates as spatial features, with a weighting factor of 2). LOCATE performance was tested against the manual segmentation using the metrics described in section 2.5. Leave-one-subject-out testing was carried out independently on the two datasets. In order to evaluate LOCATE performance in lesions with different characteristics, we additionally calculated the performance metrics in periventricular and deep lesions separately. We adopted the 10 mm distance rule: clusters within 10 mm distance from the ventricles were considered as periventricular lesions, otherwise as deep lesions (DeCarli et al., 2005). While there are other criteria available for identifying deep and periventricular lesions, the 10 mm distance rule is more suitable for automatic implementation, it is commonly used and agrees with the human rater, even in most of the confluent lesion cases (Griffanti et al., 2017). We also compared the results obtained with LOCATE with respect to the use of the optimal global threshold using the metrics described in section 2.5.

Finally, we used CADASIL and HC datasets to further validate the robustness of LOCATE with respect to lesion load and the flexibility of LOCATE with respect to the training dataset. Since the CADASIL subjects have extremely high lesion loads, while HC have negligible lesion loads, we used these datasets to test LOCATE performance in two extreme circumstances. Moreover, since manual segmentation was not available for these datasets, we used the OXVASC dataset for training (both for obtaining the lesion probability map with KNN using the same features/options, and for LOCATE) since the images were acquired using the same MRI protocol. The output obtained with LOCATE on these datasets was qualitatively evaluated and quantitatively compared with the lesion mask obtained by applying the optimal global threshold (0.9), determined with leave-one-out on the training dataset. In addition, on the CADASIL subjects, we compared LOCATE output with the mask obtained by applying a lower global threshold (0.2), which was empirically chosen (based on visual inspection of the lesion probability map) by an expert neurologist [CLH] as the optimal global threshold for this specific dataset.

### 2.5. Performance evaluation metrics

We evaluated the lesion segmentation results using the following overlap measures and detection rates:

- Dice Similarity Index (SI): calculated as 2 × (true positive lesion voxels) / (true lesion voxels + positive voxels). True lesion voxels refer to the lesion voxels in the manual segmentation and positive lesion voxels are the voxels labelled as lesions by the classifier. We used SI to generate SI plots at different thresholds and to perform paired t-tests between existing BIANCA and all the options used in the experiments described in the above sections.
- True positive rate (TPR): number of true positive lesion voxels divided by the number of true lesion voxels
- False positive rate (FPR): number of false positive lesion voxels (voxels incorrectly labelled as lesion) divided by the number of non-lesion voxels. We used TPR and FPR in order to plot ROC curves for evaluating the segmentation of existing BIANCA and all the options used in the experiments.
- Cluster-level true positive rates in deep and periventricular white matter: number of true positive lesions divided by total number of true lesions, calculated separately for deep and periventricular lesions. We used this metric to evaluate the robustness of LOCATE and to perform paired t-test between the global threshold and LOCATE results.

## 3. Results

### 3.1 Use of population-level parametric lesion probability map (PPLPM)

Figure 5 shows the ROC curves and SI plots at different thresholds for our experiments using the PPLPM with respect to age on the OPTIMA dataset. Using PPLPM as an additional feature does not improve the performance of BIANCA, although the use of the age-specific 3D map (figure 5a) still gives better results than the use of an average map across all age groups (figure 5b). When the PPLPM is post-multiplied with the subject’s lesion probability map (figure 5b), the TPR and SI values are very low, especially at higher thresholds.

**Figure 5:**
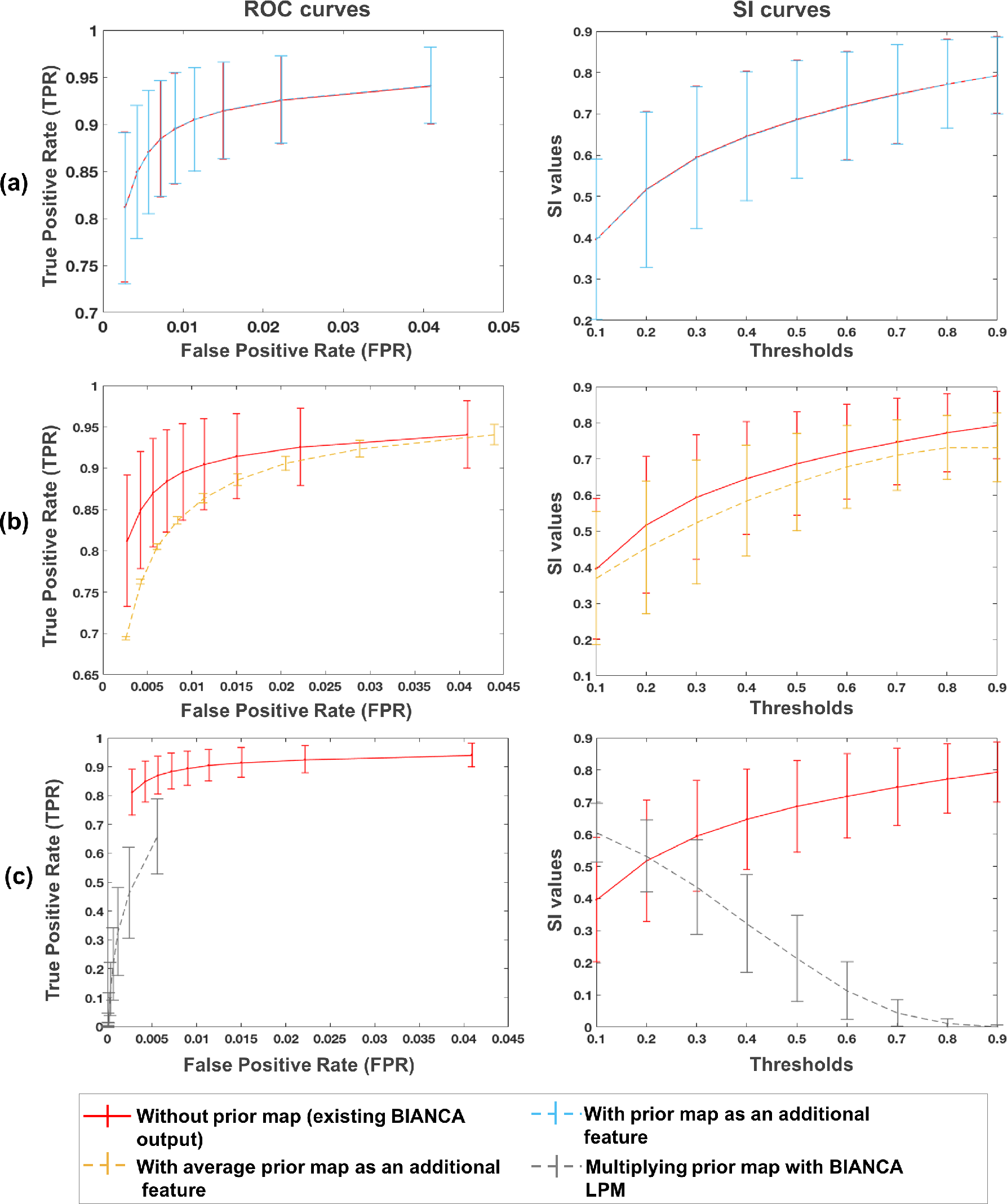
ROC (left) and SI (right) curves in the case of use of PPLPM when (a) the age-specific 3D map is added as an additional feature (light blue), (b) the average 3D map is added as an additional feature (yellow), and (c) when age-specific 3D map is post-multiplied with BIANCA lesion probability map (grey). The curves corresponding to the existing BIANCA results (Griffanti et al., 2016) are shown in red for comparison in all the three cases.

The optimal threshold for the existing version of BIANCA for this dataset was found to be 0.9 (Griffanti et al., 2016). After adding the age-specific 3D map as a new feature, we still found 0.9 to be the optimal threshold for both age-specific and average cases, while for the post-multiplication case the optimal threshold was 0.1. The paired t-test showed significantly higher SI values of existing BIANCA (at the threshold of 0.9) compared to all cases using the parametric map (p = 0.02 for age-specific map as feature, threshold 0.9; p = 0.002 for average map as feature, threshold 0.9; *p*<0.001 for the post-multiplication, threshold 0.1).

### 3.2 Comparison of alternative classifiers within BIANCA

Table 2 lists the best performing set of parameters that were obtained for each classifier on the four subjects selected from the OPTIMA dataset in the optimisation phase, when we tested using the area under the curve values for all possible combinations of the parameter values listed in table 1.

**Table 1:** Parameters considered for tuning different classifiers

| Classifier | Available main parameters and default values | Values of parameters considered for tuning |
| --- | --- | --- |
| Random forest (RF) | No. of trees - 10<br>Criteria for splitting - MSE<br>Maximum depth of trees - Till the leaves are pure<br>Minimum no. of samples required for a split - 2<br>Minimum samples at each leaf node - 1<br>Maximum number of features for each tree - No. of features<br>Minimum impurity measure at each split - $1 \times 10^{-7}$<br>Bootstrapping done - True<br>Out-of-bag error score - False | No. of trees - 20, 30, 40, 50<br>Minimum samples at each leaf node - 200, 400, 500, 600, 800, 1000<br>Minimum impurity measure at each split - $1 \times 10^{-7}$, $1 \times 10^{-6}$, $1 \times 10^{-5}$, $1 \times 10^{-4}$, $1 \times 10^{-3}$, $1 \times 10^{-2}$<br>Maximum depth of trees - 25, 50, 100 |
| Neural network (NN) | No. of hidden layers - 100<br>Activation function - RELU<br>Solver - ADAM<br>($\beta_1 = 0.9, \beta_2 = 0.999, \epsilon = 1 \times 10^{-8}$)<br>Alpha - $1 \times 10^{-4}$<br>Change in learning rate - Constant<br>Initial learning rate - $1 \times 10^{-3}$<br>Maximum no. of iterations - 200<br>Tolerance - $1 \times 10^{-4}$<br>Momentum - 0.9<br>Fraction of samples for validation - 0.1 | No. of hidden layers - 100, 120, 150<br>Solver - LBFGS, ADAM<br>Initial learning rate - $1 \times 10^{-4}$, $1 \times 10^{-3}$<br>Maximum no. of iterations - 1000, 1500, 1800<br>Tolerance - $1 \times 10^{-4}$, $1 \times 10^{-3}$, $1 \times 10^{-2}$ |
| Support vector machine (SVM) | Type of kernel - Radial basis function (RBF)<br>$\gamma$ - $1/(\text{no. of features})$<br>Tolerance - $1 \times 10^{-3}$<br>C-value - 1.0<br>Epsilon - 0.1<br>Maximum no. of iterations - Until convergence | $\gamma$ - 0.001, 0.01, 0.1, 1.0, 10<br>C-value - 0.1, 1.0, 10<br>Maximum no. of iterations - 1000, 2000, 3000, 5000, 6000 |
| Adaboost classifier (AB) | Base estimator - Decision Tree Regressor<br>No. of trees in base estimator - 1<br>No. of estimators - 50<br>Learning rate - 1.0<br>Loss function - Linear | No. of trees in base estimator - 2<br>No. of estimators - 30, 40, 50<br>Learning rate - 0.5, 0.75, 1<br>Loss function - linear, exponential, square |

**Table 2:** Best performing set of parameters for different classifiers

| Classifier | Best performing set of parameters |
| --- | --- |
| Random forest (RF) | No. of trees - 50<br>Min. samples at each leaf node - 400<br>Min. impurity measure at each split - $1 \times 10^{-3}$<br>Max. depth of trees - 25 |
| Neural network (NN) | No. of hidden layers - 100<br>Solver - LBFGS, Initial learning rate - $1 \times 10^{-3}$<br>Max. no. of iterations - 1800<br>Tolerance - $1 \times 10^{-3}$ |
| Support vector machine (SVM) | $\gamma$ - 1.0<br>C-value - 1.0<br>Max. no. of iterations - 5000 |
| Adaboost classifier (AB) | No. of trees in base estimator - 2<br>No. of estimators - 30<br>Learning rate - 1.0<br>Loss function - exponential |

Figure 6 shows the ROC curves and SI plots for the four classifiers (RF, NN, SVM, AB) using the best and the default set of parameters, with respect to KNN, on the remaining 17 subjects from the OPTIMA dataset. From the ROC curves and SI indices, overall KNN performs better than other classifiers, even for their best set of parameters. While the optimal threshold for the lesion probability map obtained from KNN classifier is 0.9, with maximum SI of 0.77, figure 6 shows that RF, NN, SVM and AB gave the maximum SI values of 0.76, 0.75, 0.65, 0.75 at thresholds 0.9, 0.8, 0.3 and 0.7 respectively. The paired t-test results, using individual subject segmentations, showed that SI values from KNN (threshold 0.9) are significantly higher than those of RF (*p* = 0.02 at threshold 0.9), SVM (*p* = 1.2 × 10^−4^ at threshold 0.3) and AB (*p* = 9.11 × 10^−4^ at threshold 0.7), while SI values using NN are not significantly different from KNN (*p* = 0.32 at threshold 0.8).

**Figure 6:**
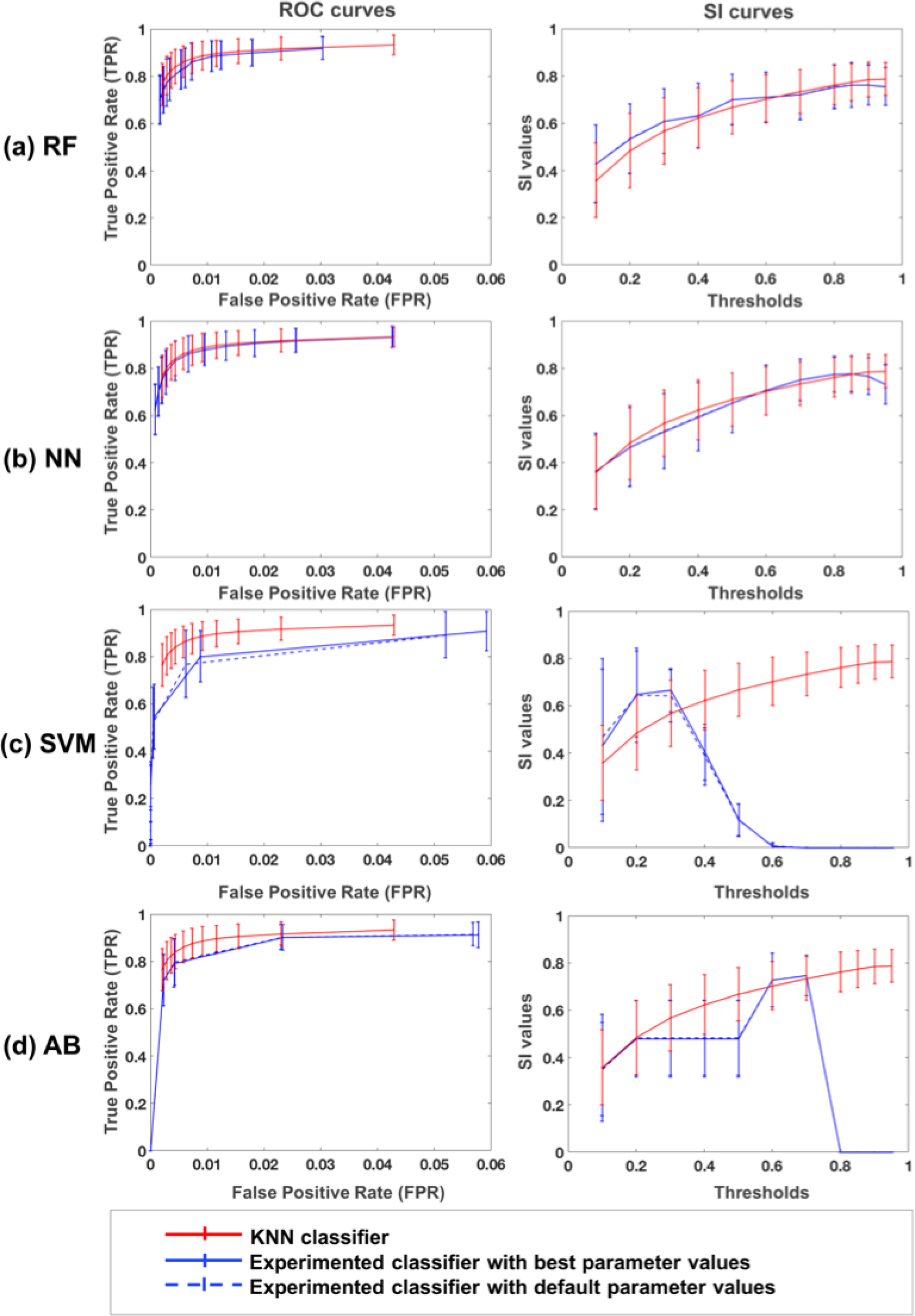
ROC curves (left) and SI curves (right) for alternative classifiers: (a) random forest (RF), (b) neural network (NN), (c) support vector machine (SVM) and (d) adaboost classifier (AB). Results are shown for each classifier’s best (blue solid line) and default parameters (blue dashed lines) along with the results for KNN classifier, currently used in BI1A5NCA (red solid line).

### 3.3 LOCally Adaptive Threshold Estimation (LOCATE)

Figure 7 illustrates examples of BIANCA results with LOCATE outputs compared to the global thresholded outputs and the manual segmentation: LOCATE detected more deep lesions (Fig. 7c, d) and segmented periventricular lesions better (Fig. 7a, b, d) with respect to the manual segmentation.

**Figure 7:**
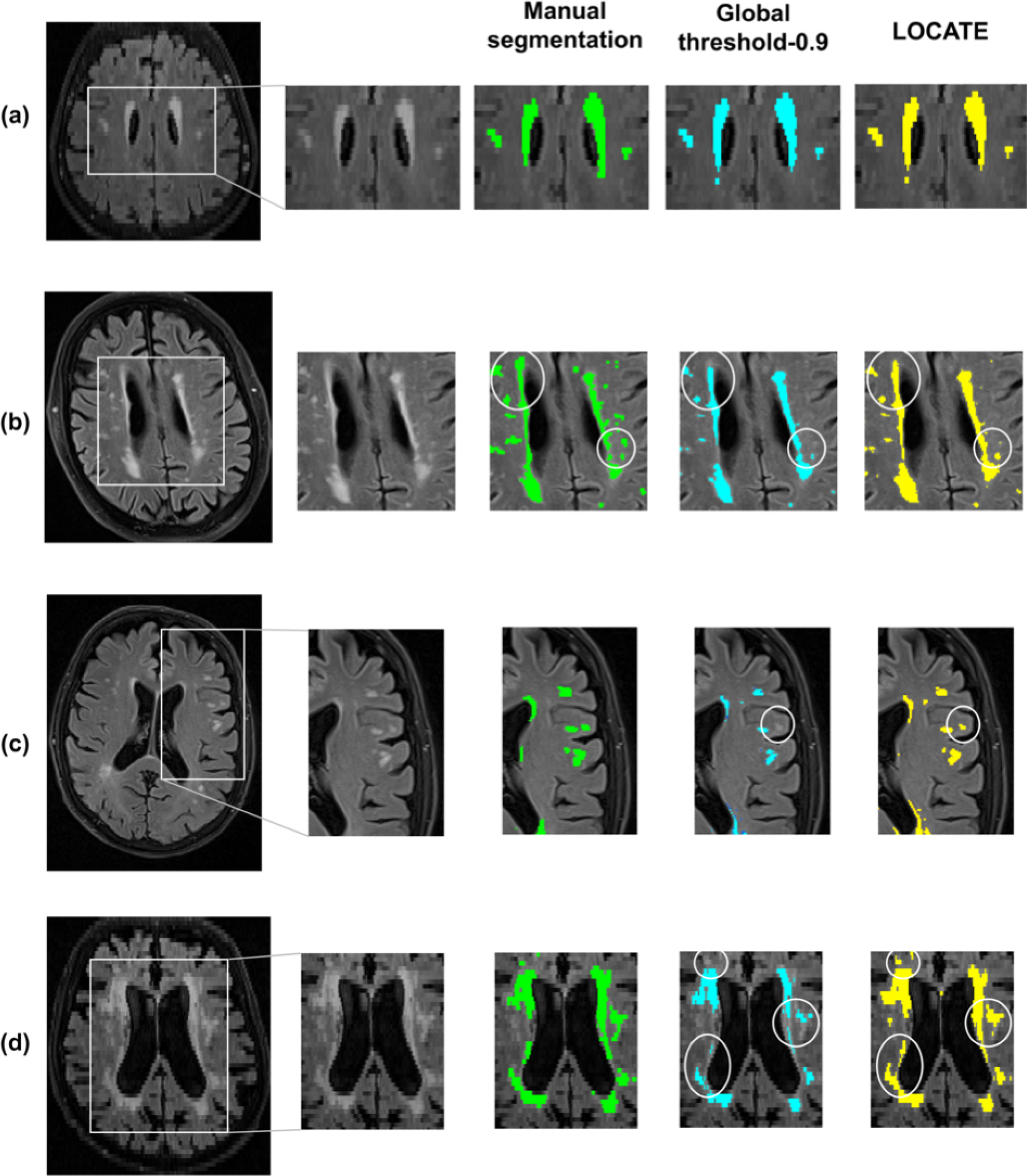
Example results of LOCATE for (a) low, (b) medium, (c) high and (d) very high lesion load (a,d from the OXVASC; b,c from the OPTIMA datasets). Manual segmentation (green) is shown along with outputs of global thresholding at 0.9 (light blue) and LOCATE (yellow). LOCATE provides better segmentation in deep (c, d) and periventricular white matter lesions (a, b, d) when compared to global thresholding.

Figure 8 shows the ROC curves, SI plots and cluster-wise true positive rate in periventricular and deep white matter for BIANCA at various global thresholds and using LOCATE for OPTIMA and OXVASC datasets. The SI values obtained with LOCATE were not significantly different according to the paired t-test to those obtained with global thresholding values of 0.9 both for the OPTIMA (LOCATE SI = 0.77 ± 0.10; global threshold SI = 0.77 ± 0.08, *p* = 0.94) and for the OXVASC (LOCATE SI = 0.75 ± 0.14; global threshold SI = 0.74 ± 0.11, *p* = 0.94) datasets. However, in terms of detection rates, LOCATE gave an increase in voxel-wise TPR (0.03 for the OPTIMA and 0.10 for the OXVASC) with a negligible increase in FPR (0.001 for the OPTIMA and 0.002 for the OXVASC) compared to the global threshold of 0.9.

**Figure 8:**
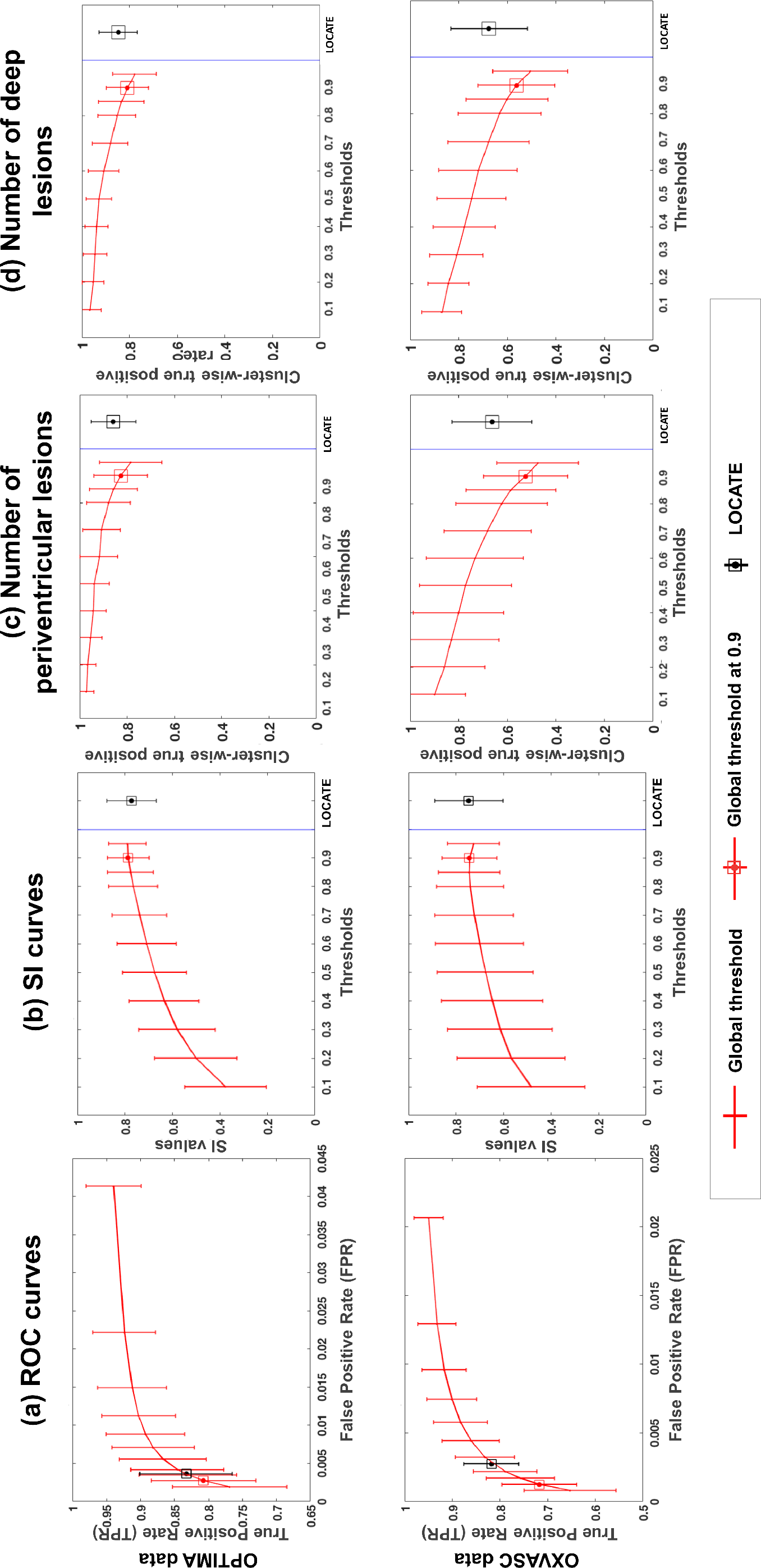
Comparison of ROC curves, SI plots and true positive lesion counts in periventricular and deep region for LOCATE and global thresholding. ROC curves (a), SI indices (b) and true positive lesion counts in periventricular (c) and deep (d) regions shown for the OPTIMA (top row) and the OXVASC (bottom row) datasets.

The results of the paired t-test show that the cluster-wise true positive rates for LOCATE are significantly higher than those for global thresholding in both periventricular white matter (figure 8c, *p* = 0.003 for the OPTIMA and *p*<0.001 for the OXVASC) and deep white matter (figure 8d, *p*<0.001 for the OPTIMA and *p*<0.001 for the OXVASC).

Figure 9 illustrates the results of lesion segmentation with LOCATE in four subjects from the CADASIL dataset. Since CADASIL patients have different lesion characteristics compared to those in the OXVASC dataset (different lesion location and very high lesion loads), applying the global threshold of 0.9 that was determined in the training phase on the OXVASC dataset yields poorly segmented binary lesion maps (figure 9b). The global threshold visually determined as optimal (by an expert neurologist CLH) was 0.2 (figure 9c), however this leads to increased false positives in some cases, as shown in the third case of figure 9c. LOCATE provides a much better segmentation and detects the lesions in the temporal lobe, typical of the pathology (figure 9d). A similar comparison of LOCATE results with those of global thresholding at 0.9 on the HC dataset is shown in figure 10.

**Figure 9:**
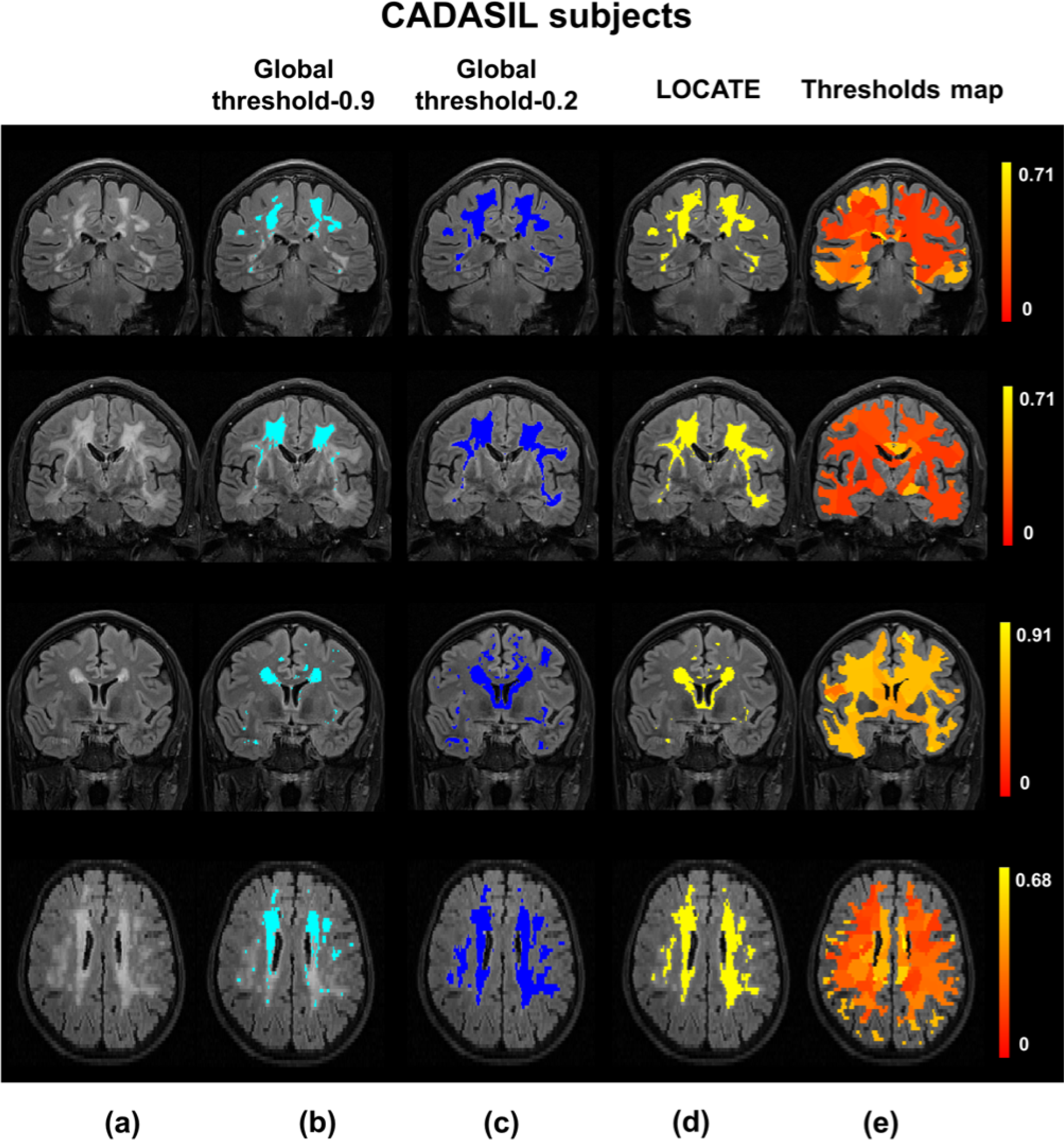
Example results of LOCATE on the CADASIL data. (a) FLAIR image, (b) global thresholding at 0.9 (light blue), (c) global thresholding at 0.2 (dark blue), (d) LOCATE results (yellow) and (e) threshold maps obtained from LOCATE (red-yellow). Note that the threshold maps shows the heterogeneity in the lesions probabilities in various regions of the white matter.

**Figure 10:**
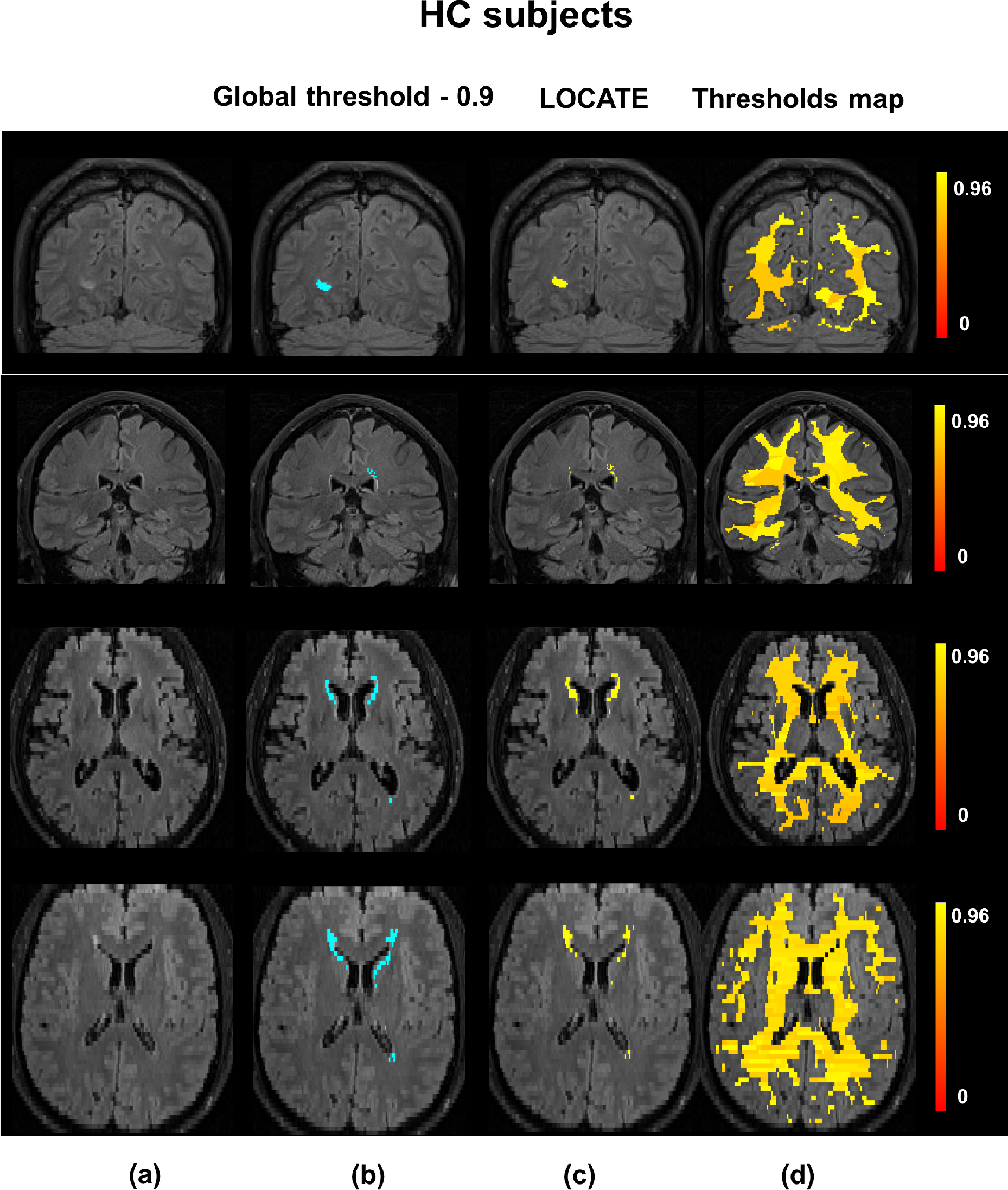
Example results of LOCATE on the HC data. (a) FLAIR image, (b) global thresholding at 0.9 (light blue), (c) LOCATE results (yellow) and (d) threshold maps obtained from LOCATE (red-yellow). Note that the threshold maps shows the heterogeneity in the lesions probabilities in various regions of white matter.

When the binary lesion maps obtained using the optimal global threshold value of 0.2 is used as the reference segmentation, the paired t-test results show that the SI values obtained for CADASIL with LOCATE (SI = 0.79 ± 0.01) are significantly higher than those obtained using a global threshold at 0.9 (SI = 0.57 ± 0.10, *p*<0.001). For CADASIL, using LOCATE increases the voxel-wise TPR by 0.48 for a constant FPR of 0.00.

To summarise the effect of LOCATE on different datasets, figure 11 shows boxplots of local threshold values determined by LOCATE for the four datasets, along with the optimal global threshold values for each dataset. The median threshold values of OPTIMA and HC datasets are higher than other two datasets, especially the CADASIL dataset. LOCATE assigned higher thresholds for datasets with lower lesion load (HC and OPTIMA, which contains 30% of healthy subjects) and lower threshold for datasets with higher lesion load (OXVASC and especially CADASIL). This is also observable from the thresholds maps shown in figure 9e and figure 10e.

**Figure 11:**
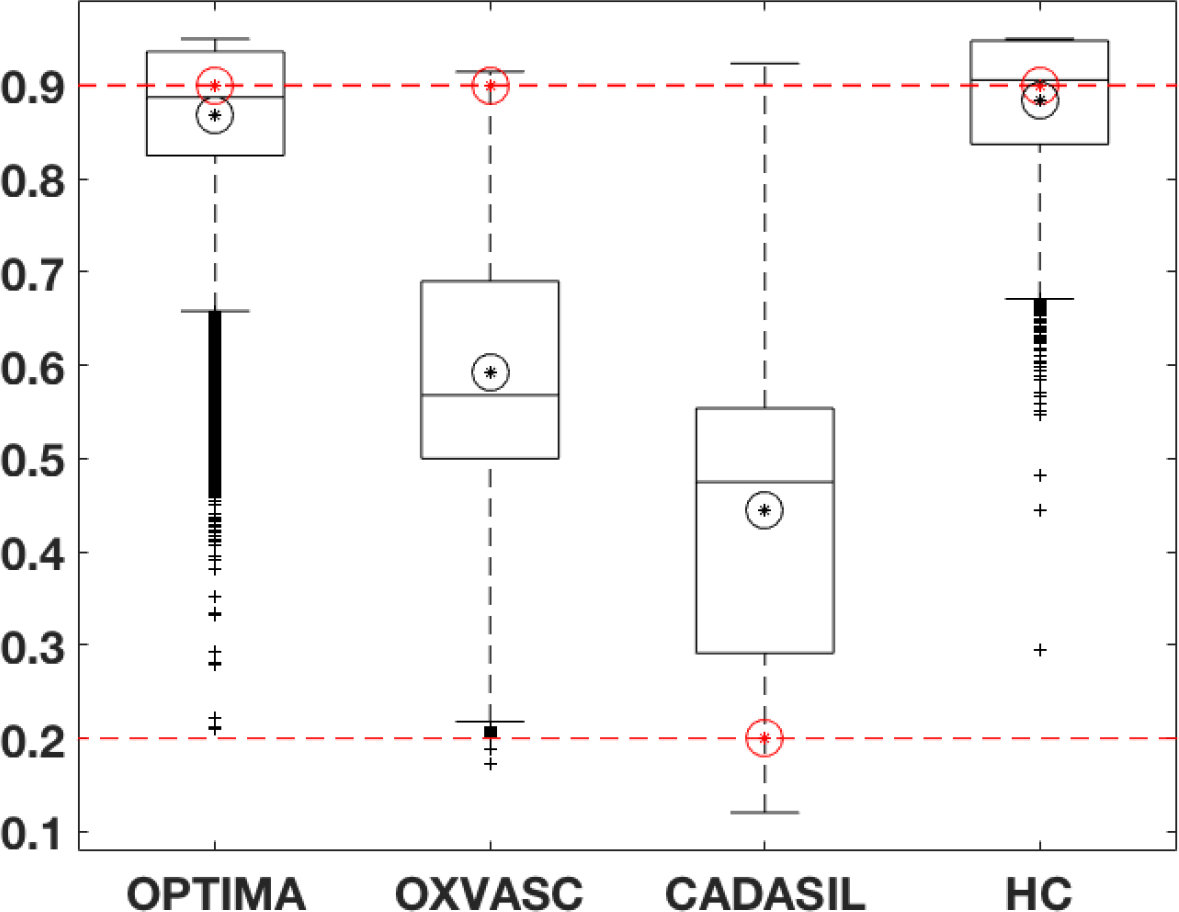
Boxplots of local thresholds obtained from the random forest regression model in LOCATE. The optimal global thresholds determined manually are shown in red circles. The mean values of local thresholds obtained from LOCATE are shown in black circles.

In line with the results of the paired t-test, the largest improvement in SI corresponds to the largest difference between the LOCATE threshold and the global threshold. In the case of the HC dataset, the median of local thresholds obtained from LOCATE is very close to the optimal threshold of 0.9, thus showing only a slight improvement in the lesion segmentation. This is also evident from figure 10, where LOCATE results appear very similar to the results of global thresholding at 0.9 (figure 10b and 10c).

## 4. Discussion

In this work, we studied the effect of single-subject and population-level heterogeneity in amount, location and characteristics of WMH on our recently developed WMH segmentation method BIANCA, with the aim of improving both the classification and the thresholding steps. At the classification step, we analysed the effect on the subject-level lesion probability maps of using PPLPM (with age as factor of interest) and of using alternative classifiers, that have been previously used for this task in the literature. For thresholding the lesion probability map, as an alternative to selecting a global threshold, we proposed LOCATE, a method to determine local thresholds that are sensitive to spatial differences in the lesion probabilities, giving more accurate binary lesion masks.

We observed the effect of using the PPLPM and found that post-multiplying the age-specific 3D map with the lesion probability map performs worse than adding the PPLPM as an additional feature. This is because PPLPM indicates the likelihood of finding a lesion at a given location in a population, informed primarily by the anatomy of disease signs. On the other hand, the subject-level probability map reflects the probability of identifying a lesion versus the normal tissue and is primarily determined by the intensity contrasts and noise in the image. For instance, in our previous work (Sundaresan et al., 2018, *Preprint, under review*) we showed that in PPLPM the lesion probabilities typically increase more in the periventricular WM than in the deep WM in the elder population (which is in line with the existing literature Simoni et al., 2012; Van Den Heuvel et al., 2004; Sabayan et al., 2015). Therefore, post-multiplying the above PPLPM with the subject-level lesion probability map can bias the subject-level lesion probabilities more towards the periventricular lesions, irrespective of the local spatial characteristics in the images from the individual subject. However, when we provide the PPLPM as an additional feature, the classifier assigns sensible lesion probability values taking into account the PPLPM and other additional features such as image intensity contrasts for the different modalities and spatial coordinates. Particularly, we observed that adding age-specific map as an additional feature gives better results than the average map alone, since the former models the lesion probabilities for the subject’s age group specifically and hence is more accurate. Overall we observed that, at present, using the PPLPM does not improve significantly the performance of BIANCA (also evident from the p-values of our t-test results). However, the way this option will be implemented in a future release of BIANCA will allow the user to use any 4D PPLPM with respect to any other factor of interest, which could improve the segmentation in their specific dataset.

The results on the comparison between KNN and other classifiers showed that KNN provided better segmentation. For the same set of features (excluding the PPLPM), we observed that SVM and AB classifiers provided significantly lower SI than KNN, especially at higher thresholds. On the other hand, SI values for RF and NN followed a similar trend to KNN with relatively similar SI values (figure 6). SI values for RF were still significantly lower than KNN, while NN gave SI values that were not significantly different from KNN. Moreover, for a given FPR, KNN has a higher TPR compared to any other classifier, indicating that KNN detects more true lesions than other classifiers. However, since NN and RF have the potential to perform as well as KNN, they will be included as alternative classifiers in a future release of BIANCA. At this point, it is worth noting that irrespective of the classifier used, there is a certain degree of uncertainty associated with the manual segmentation – they are an approximation to the real ground truth, meaning that very fine discrimination may reflect errors in the manual segmentation rather than errors in the automated classification results.

Regarding the improvements at the thresholding step with LOCATE, the initial leave-one-subject-out results of LOCATE on the OPTIMA and the OXVASC datasets showed improvement in BIANCA segmentation for both the datasets. LOCATE detected more true positive lesions, especially in the deep white matter, and provided better delineation of the lesions with respect to the manual segmentation as indicated by the improved SI values and the visual results shown in figure 7. Our validation of LOCATE on CADASIL and HC datasets proves that LOCATE is robust with respect to the extreme variations in lesion load and location without the need to retrain BIANCA. In fact, LOCATE can be trained on any data having the same modalities, acquired with the same sequence to the test dataset. For instance, we used the OXVASC dataset to train LOCATE for testing it on the CADASIL dataset since both datasets were acquired with the same sequence.

The pattern of lesions for CADASIL subjects differs from that of vascular subjects in the OXVASC dataset, as CADASIL subjects, in addition to the normal distribution of WMH seen in SVD, will typically have lesions within the external capsule and anterior temporal lobes (Chabriat et al., 2009). Due to this variability in lesion pattern, training BIANCA on the OXVASC dataset using features that include spatial coordinates yields very low lesion probabilities in the temporal regions. Reflecting these lower lesion probabilities, the thresholds map from LOCATE (figure 9) shows much lower threshold values, especially in the temporal regions in the CADASIL images. Therefore LOCATE yields much better segmentation performance (SI = 0.79) compared to a global threshold of 0.9 (SI = 0.57), when the lesion probability map thresholded at 0.2 was considered as the reference segmentation. Moreover, LOCATE results on CADASIL showed a substantial increase of 0.48 in TPR without an increase in FPR, due to the detection of more lesions in the temporal region. This indicates that LOCATE takes into account different lesion patterns and hence can be used in different pathologies.

The threshold maps obtained from LOCATE for the HC dataset show higher thresholds when compared with the CADASIL dataset. This is due to the fact that HC have mostly periventricular lesions (that are common in healthy ageing) that are usually assigned higher probabilities. Hence by using higher threshold values, LOCATE gives results that appear visually similar to the global thresholding at 0.9, detecting mostly periventricular lesions. While the threshold maps from LOCATE (figure 9 and 10) provide information regarding the spatial heterogeneity of lesion probabilities, boxplots of thresholds shown in figure 11 show the overall characteristics of the datasets. The improvement in segmentation performance is more prominent in OXVASC and CADASIL datasets when compared to OPTIMA and HC datasets. This is reflected in their corresponding optimal global thresholds being very different from the median of their LOCATE thresholds, which are more adaptive to spatially local lesion probabilities in deep and periventricular white matter. LOCATE results on the four datasets, especially the HC dataset (which has negligible lesion load compared with the CADASIL subjects) show that LOCATE is more adaptive to the variation in the global lesion load in addition to the spatial heterogeneity in lesion probabilities, compared to global thresholding (shown in figure 9 and 10).

LOCATE can be used when manual segmentation is unavailable for the test dataset, or the number of subjects in the dataset is not enough to generate an accurate training dataset. In its current Matlab implementation, LOCATE running time to determine a binary lesion map from an individual lesion probability map is approximately 15 minutes per subject, when run on an iMac with a 2.9 GHz Intel Core i5processor.

LOCATE will be included as an additional option for generating binary lesion maps, as an alternative to the faster, but less accurate, global thresholding, in a future release of BIANCA.

## Acknowledgements

We acknowledge all the participants. We are grateful to Prof. Gordon K. Wilcock and all the staff of OPTIMA. We acknowledge Prof Hugh Markus and Asst. Prof Andrea Nemeth for their help in recruiting the CADASIL cohort. We acknowledge the use of the facilities of the Acute Vascular Imaging Centre, Oxford. We also thank Dr. Chiara Vincenzi and Dr. Francesco Carletti for their help on generating the manual masks used in our experiments.

M.J. receives royalties from licensing of FSL to non-academic, commercial parties. The authors report no potential conflicts of interest.

## References

Anbeek, P., Vincken, K.L., Van Osch, M.J., Bisschops, R.H., Van Der Grond, J.. Probabilistic segmentation of white matter lesions in MR imaging. NeuroImage 2004;21(3):1037–1044.

Attems, J., Jellinger, K.A.. The overlap between vascular disease and alzheimers disease-lessons from pathology. BMC medicine 2014;12(1):206.

Breiman, L.. Random forests. Machine learning 2001;45(1):5–32.

Chabriat, H., Joutel, A., Dichgans, M., Tournier-Lasserve, E., Bousser, M.G.. Cadasil. The Lancet Neurology 2009;8(7):643–653.

Charlton, R.A., Morris, R., Nitkunan, A., Markus, H.. The cognitive profiles of cadasil and sporadic small vessel disease. Neurology 2006;66(10):1523–1526.

De Boer, R., Vrooman, H.A., Van Der Lijn, F., Vernooij, M.W., Ikram, M.A., Van Der Lugt, A., Breteler, M.M., Niessen, W.J.. White matter lesion extension to automatic brain tissue segmentation on MRI. NeuroImage 2009;45(4):1151–1161.

Debette, S., Markus, H.. The clinical importance of white matter hyperintensities on brain magnetic resonance imaging: systematic review and meta-analysis. BMJ 2010;341:c3666.

DeCarli, C., Fletcher, E., Ramey, V., Harvey, D., Jagust, W.J.. Anatomical mapping of white matter hyperintensities (wmh): exploring the relationships between periventricular wmh, deep wmh, and total wmh burden. Stroke 2005;36(1):50–55.

Dyrby, T.B., Rostrup, E., Baaré, W.F., van Straaten, E.C., Barkhof, F., Vrenken, H., Ropele, S., Schmidt, R., Erkinjuntti, T., Wahlund, L.O., et al. Segmentation of age-related white matter changes in a clinical multi-center study. NeuroImage 2008;41(2):335–345.

Geremia, E., Clatz, O., Menze, B.H., Konukoglu, E., Criminisi, A., Ayache, N.. Spatial decision forests for MS lesion segmentation in multi-channel magnetic resonance images. NeuroImage 2011;57(2):378–390.

Griffanti, L., Jenkinson, M., Suri, S., Zsoldos, E., Mahmood, A., Filippini, N., Sexton, C.E., Topiwala, A., Allan, C., Kivimäki, M., et al. Classification and characterization of periventricular and deep white matter hyperintensities on mri: A study in older adults. NeuroImage 2017;.

Griffanti, L., Zamboni, G., Khan, A., Li, L., Bonifacio, G., Schulz, U.G., Kuker, W., Battaglini, M., Rothwell, P.M., Jenkinson, M.. BIANCA (brain intensity abnormality classification algorithm): a new tool for automated segmentation of white matter hyperintensities. NeuroImage 2016;.

Hernändez, M.D.C.V., Chappell, F.M., Maniega, S.M., Dickie, D.A., Royle, N.A., Morris, Z., Anblagan, D., Sakka, E., Armitage, P.A., Bastin, M.E., et al. Metric to quantify white matter damage on brain magnetic resonance images. Neuroradiology 2017;59(10):951–962.

Kim, K.W., MacFall, J.R., Payne, M.E.. Classification of white matter lesions on magnetic resonance imaging in elderly persons. Biological psychiatry 2008;64(4):273–280.

Le Heron, C., Manohar, S., Plant, O., Muhammed, K., Griffanti, L., Nemeth, A., Douaud, G., Markus, H., Husain, M.. Dysfunctional effort-based decision making underlies apathy in genetic cerebral small vessel disease. BRAIN 2018;.

Li, L., Simoni, M., Küker, W., Schulz, U.G., Christie, S., Wilcock, G.K., Rothwell, P.M.. Population-based case–control study of white matter changes on brain imaging in transient ischemic attack and ischemic stroke. Stroke 2013;44(11):3063–3070.

Mitra, J., Bourgeat, P., Fripp, J., Ghose, S., Rose, S., Salvado, O., Connelly, A., Campbell, B., Palmer, S., Sharma, G., et al. Lesion segmentation from multimodal MRI using random forest following ischemic stroke. NeuroImage 2014;98:324–335.

Pantoni, L.. Cerebral small vessel disease: from pathogenesis and clinical characteristics to therapeutic challenges. The Lancet Neurology 2010;9(7):689–701.

Rostrup, E., Gouw, A., Vrenken, H., van Straaten, E.C., Ropele, S., Pantoni, L., Inzitari, D., Barkhof, F., Waldemar, G., Group, L.S., et al. The spatial distribution of age-related white matter changes as a function of vascular risk factorsresults from the ladis study. Neuroimage 2012;60(3):1597–1607.

Rothwell, P., Coull, A., Giles, M., Howard, S., Silver, L., Bull, L., Gutnikov, S., Edwards, P., Mant, D., Sackley, C., et al. Change in stroke incidence, mortality, case-fatality, severity, and risk factors in oxfordshire, UK from 1981 to 2004 (oxford vascular study). The Lancet 2004;363(9425):1925–1933.

Sabayan, B., van der Grond, J., Westendorp, R.G., van Buchem, M.A., de Craen, A.J.. Accelerated progression of white matter hyperintensities and subsequent risk of mortality: a 12-year follow-up study. Neurobiology of aging 2015;36(6):2130–2135.

Simoni, M., Li, L., Paul, N.L., Gruter, B.E., Schulz, U.G., Küker, W., Rothwell, P.M.. Age- and sex-specific rates of leukoaraiosis in TIA and stroke patients population-based study. Neurology 2012;79(12):1215–1222.

Sundaresan, V., Griffanti, L., Kindalova, P., Alfaro-Almagro, F., Zamboni, G., Rothwell, P.M., Nichols, T.E., Jenkinson, M.. Modelling the distribution of white matter hyperintensities due to ageing on mri images using bayesian inference. bioRxiv 2018;.

Van Den Heuvel, D., Admiraal-Behloul, F., Ten Dam, V., Olofsen, H., Bollen, E., Murray, H., Blauw, G., Westendorp, R., De Craen, A., Van Buchem, M., et al. Different progression rates for deep white matter hyperintensities in elderly men and women. Neurology 2004;63(9):1699–1701.

Vannorsdall, T.D., Waldstein, S.R., Kraut, M., Pearlson, G.D., Schretlen, D.J.. White matter abnormalities and cognition in a community sample. Archives of clinical neuropsychology 2009;24(3):209–217.

Wardlaw, J.M., Smith, E.E., Biessels, G.J., Cordonnier, C., Fazekas, F., Frayne, R., Lindley, R.I., T O’Brien, J., Barkhof, F., Benavente, O.R., et al. Neuroimaging standards for research into small vessel disease and its contribution to ageing and neurodegeneration. The Lancet Neurology 2013;12(8):822–838.

Wen, W., Sachdev, P.. The topography of white matter hyperintensities on brain mri in healthy 60-to 64-year-old individuals. Neuroimage 2004;22(1):144–154.

Yamamoto, D., Arimura, H., Kakeda, S., Magome, T., Yamashita, Y., Toyofuku, F., Ohki, M., Higashida, Y., Korogi, Y.. Computer-aided detection of multiple sclerosis lesions in brain magnetic resonance images: False positive reduction scheme consisted of rule-based, level set method, and support vector machine. Computerized Medical Imaging and Graphics 2010;34(5):404–413.

Zamboni, G., Griffanti, L., Jenkinson, M., Mazzucco, S., Li, L., Küker, W., Pendlebury, S.T., Rothwell, P.M.. White matter imaging correlates of early cognitive impairment detected by the montreal cognitive assessment after transient ischemic attack and minor stroke. Stroke 2017;48(6):1539–1547.

Zamboni, G., Wilcock, G.K., Douaud, G., Drazich, E., McCulloch, E., Filippini, N., Tracey, I., Brooks, J.C., Smith, S.M., Jenkinson, M., et al. Resting functional connectivity reveals residual functional activity in alzheimers disease. Biological psychiatry 2013;74(5):375–383.

